# A Transcriptomic Comparison of the HD10.6 Human Sensory Neuron-Derived Cell Line with Primary and iPSC Sensory Neurons

**DOI:** 10.1101/2025.04.03.643725

**Authors:** Zaid Al-Abbasi, Shamsuddin A. Bhuiyan, William Renthal, Derek C. Molliver

## Abstract

A key concern in early-stage analgesic discovery efforts is the extent to which mechanisms identified in rodents will translate to humans. To evaluate an alternative approach to the use of rodent dissociated DRG neurons for in vitro analyses of nociceptive signaling, we performed a transcriptomic analysis of the HD10.6 human dorsal root ganglion (DRG)-derived immortalized cell line. We conducted RNA-seq on proliferating and mature HD10.6 cells to characterize transcriptional changes associated with maturation. We then compared the transcriptomes of HD10.6 cells and several recently developed lines of human induced pluripotent stem cell-derived sensory neurons (iPSC-SN) to single-nucleus RNA-seq data from human DRGs. HD10.6 cells showed the highest correlation with 3 human sensory neuron subtypes associated with nociception and pruriception. Each of the iPSC-SN lines evaluated showed a distinct pattern of correlation with human sensory neuron subtypes. We identified G protein-coupled receptors (GPCRs) and ion channels that are expressed in both HD10.6 cells and human DRG neurons, as well as numerous genes that are expressed in human DRG but not in rodent, underscoring the need for human sensory neuron in vitro models. Proof-of-concept evaluations of protein kinase A, protein kinase C and Erk signaling provide examples of scalable assays using HD10.6 cells to investigate well-established GPCR signaling pathways. We conclude that HD10.6 cells provide a versatile model for exploring human neuronal signaling mechanisms.

## INTRODUCTION

The development of more effective, non-addictive pain therapeutics requires a deeper understanding of the molecular mechanisms that give rise to chronic pain in humans. Although rodent models remain important tools for studying nociceptive processes and the development of pain-related behavior, the expanding use of human tissue for preclinical pain research has revealed mechanistic differences between human and mouse nociception [7; 18; 114; 116]. However, acquiring large numbers of high-quality live human sensory neurons for biochemical assays remains rate-limiting and requires substantial expertise and infrastructure [73; 87; 107].

To develop accessible and translational *in vitro* models for human nociceptors, a growing number of studies use human induced pluripotent stem cells differentiated into sensory neuron-like cells (iPSC-SNs) [16]. Human iPSC-SN protocols are rapidly evolving, however most current protocols require complex and time-intensive transcriptional reprogramming to generate nociceptor-like cells, and the resulting cells may diverge substantially from primary human nociceptor gene expression profiles and physiology [4; 38; 61; 83]. In addition, commercially available iPSC-SNs are expensive and are typically sold as terminally differentiated cells, limiting their utility.

HD10.6 cells are a human DRG-derived cell line that provides a potentially useful alternative to iPSC-SNs and primary DRG culture models but has only been used in a handful of publications [11; 37; 103; 118]. It is currently the only published human sensory neuron-derived immortalized cell line [33] and was produced from male human embryonic DRG cells immortalized by transfection with a tetracycline-sensitive v-myc construct [37]. Unlike iPSCs, HD10.6 cells are restricted to a neuron-like fate. The application of maturation media containing tetracycline suppresses v-myc expression, causing cells to stop dividing, extend neurites, and develop a nociceptor-like phenotype [33]. In this study, we used a transcriptomic approach to examine the utility of HD10.6 cells as an in vitro model of human nociceptors. Bulk mRNA-seq was performed on proliferating and mature HD10.6 cells, and the results were compared to published transcriptomics data from primary human DRG neurons. We also compared published data from several recently developed iPSC-SNs to human primary DRG neurons. Initial proof-of-concept evaluations of protein kinase A (PKA), protein kinase C (PKC), and extracellular signal-regulated kinase 1/2 (ERK1/2) activity provide examples of scalable assays using HD10.6 cells to investigate well-defined GPCR signaling pathways. Our results support the conclusion that the HD10.6 cell line, used in conjunction with reference human transcriptomics data, can be a valuable addition to the preclinical research toolkit for investigating human sensory neuron signaling mechanisms.

## METHODS & MATERIALS

### Cell Culture

All work reported here was performed under protocols approved by the University of New England Institutional Biosafety Committee. HD10.6 cells (RRID: CVCL_WG82) were cultured following a published protocol [12] with minor modifications as described below. Short tandem repeat analysis confirmed the HD10.6 identity as logged in the Cellosaurus online database. Cells were confirmed to be mycoplasma-negative using a commercial kit [GeneCopoeia, MP004]. The cells were cultured in T25 flasks coated with fibronectin (Millipore, FC010) and incubated in a 5% CO2 environment at 37°C. Proliferating HD10.6 cells were grown in proliferation complete medium (PCM) containing 50% advanced Dulbecco’s modified Eagle medium (DMEM) (ThermoFisher, 10567022) and 50% nutrient mixture F-12 (Fisher Scientific, 12634-010), supplemented with 1X GlutaMAX (ThermoFisher, 35050061), 1X B-27 plus neuronal supplement (B-27 plus) (ThermoFisher, A3582801), 10 ng/ml prostaglandin E1 (Sigma-Aldrich, P5515-1MG), 0.5 ng/ml beta fibroblast growth factor (R&D Systems, 3718-FB), and 50 µg/ml geneticin G418 solution (Sigma Aldrich, 4727878001). Upon reaching approximately 70% confluency in the flasks, cells were sub-cultured in PCM on plates pre-coated with 5 µg/ml poly-D-lysine (EMD Millipore, US0157762) at a density of 15 x 10^4^ cells/well in 6-well plates. After 24 hours, cell maturation was initiated by replacing the PCM with a maturation complete medium (MCM) containing Neurobasal Plus medium (Thermo Fisher, A3582901) supplemented with 1X GlutaMAX, 1X B-27 Plus, 50 ng/ml beta nerve growth factor (β-NGF) (Peprotech, 450-01), 25 ng/ml ciliary neurotrophic factor (CNTF) (Hellobio, HB8968), glial cell-derived neurotrophic factor (GDNF) (ThermoFisher, RP-8602), 1 µg/ml tetracycline (Sigma Aldrich, T7660-5G), and 25 µM forskolin (LC Labs, F-9929). On the fourth day of maturation, the media was replaced with fresh maturation media without forskolin or trophic factors. Cells were analyzed after 7 days of maturation unless otherwise noted.

### Reverse transcription PCR (RT-PCR)

All mRNA isolation procedures were performed in an RNA hood. The hood and pipettes were cleaned with 70% ethanol an-RNA-zap and UV-irradiated for 20 minutes. Low-retention pipet tips were used throughout the process. Total RNA was isolated from lysed HD10.6 cells using the Norgen RNA extraction and isolation kit (Norgen Biotek, 17200) according to manufacturer instructions. cDNA was generated using the SuperScript^TM^ II Reverse Transcriptase (Invitrogen, 18064022). RT-PCR was conducted using 1x PCR buffer, 2 mM MgCl2, 0.2 mM dNTP, 0.25 μM primers, 1.25 U GoTaq Hot start polymerase (Promega, M5005) and 400ng of cDNA, or without cDNA for negative control samples. The amplification parameters were set as follows: initial denaturation at 94°C for 2 minutes, followed by cycles at 94°C for 30 seconds, 58-62°C for 30 seconds (depending on the optimized temperature for each set of primers (**Table 1**), and 72°C for 1 minute. This cycle was repeated 30 times, followed by a 10-minute extension at 72°C.

**Table 1.**
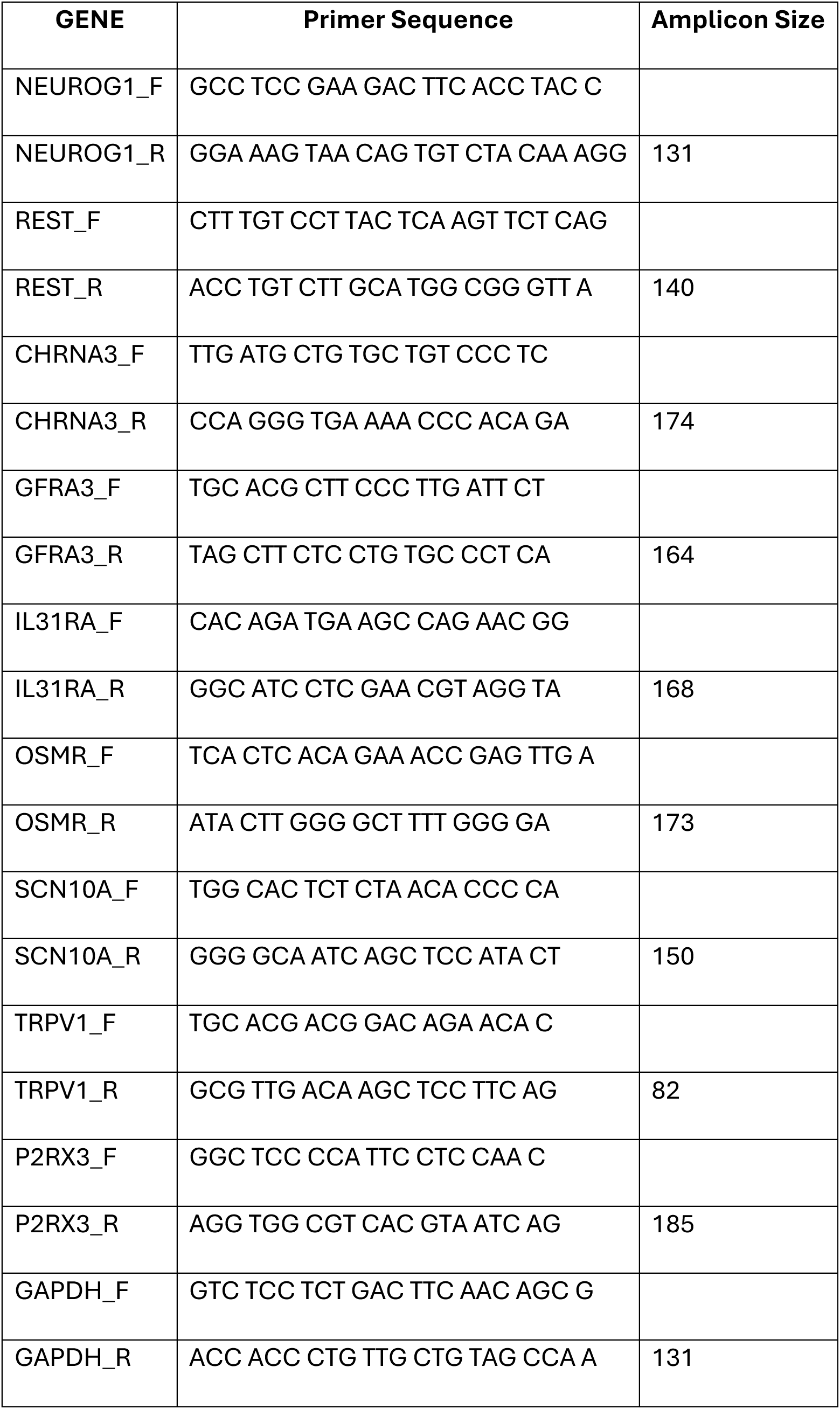
PCR primers (human).

### Quantitative PCR (qPCR)

Relative expression levels were measured by SYBR green real-time PCR (Qiagen, 330513) using the ΔCT method with glyceraldehyde 3-phosphate dehydrogenase (GAPDH) as the reference standard [74]. Additional samples lacking cDNA were used as negative controls. Twenty µl reactions were prepared as follows: 10μl SYBR green, 1μl forward and reverse primers (5μM), 1μl cDNA (400ng), and 7μl DNase/RNase-free water. The mixture was then processed for 40 cycles using the Applied Biosystems StepOnePlus™ Real-Time PCR System.

### RNA extraction for RNA-seq

RNA isolation was performed using the TrIzol Plus RNA purification kit (ThermoFisher, 12183555) according to the manufacturer’s instructions. RNA concentration was measured using a Nanodrop One (Thermo Scientific, 13-400-519), and quality was verified using an Agilent 2100 bioanalyzer. Five replicates each of proliferating and mature cells (grown for 7 days in MCM) were stored at −80°C before mRNA was isolated and shipped on dry ice to the Oklahoma Medical Research Foundation Genomics Core (Oklahoma City, OK) for next-generation sequencing using the Illumina 150 bp paired-end reads NovaSeq 6000 sequencing system.

### Bioinformatics analysis

FASTQ files were received from the genomics core, and reads were trimmed using trimmomatic [8]. We then aligned reads to the human reference genome (GRCh37) using STAR v2.7.9 and confirmed that >75% of reads aligned to the reference [19]. FeatureCounts generated counts tables were loaded into the R package DESEQ2 for transcript per million (TPM) normalization, sample similarity evaluation (PCA), and identification of differentially expressed genes between proliferating and mature HD10.6 cells [66]. For comparison of human subtype transcriptomes to previously published transcriptomes [12; 16; 92; 98; 115], data were processed as described above for our HD10.6 cells, where raw data were mapped to GRCh37 and FeatureCounts/DESEQ2 were used to generate raw and TPM-normalized counts tables. For the purposes of gene expression comparison across different bulk RNA-seq libraries, we combined raw counts for the mature HD10.6 cells and the previously published transcriptomes. We then renormalized (TPM) across the entire counts table.

To transcriptomically compare mature HD10.6 cells to defined subtypes of human DRG neurons, we correlated gene expression levels between HD10.6 cells and human subtype data from the harmonized atlas [7]. To minimize technical variations between studies, only genes identified in all datasets were included in the comparisons; approximately 30,000 genes identified in the bulk RNA-seq from HD10.6 cells were excluded, many of which were non-coding genes. Human genes in the HD10.6 counts table were converted into mouse gene names (1:1 orthologs) using Ensembl BioMart [39], and we calculated the average expression for each subtype found in the harmonized cell atlas using Seurat’s AverageExpression function for the normalized counts on only human nuclei [7; 35]. Only the human neuronal subtypes with >20 nuclei in the harmonized atlas were used for the analysis; excluded subtypes with <20 nuclei included Calca+Dcn, Rxfp1, and Th. The characterization of gene types, including ion channels and GPCRs, was obtained from the Gene Ontology provided by the PANTHER database [71].

### In vitro stimulation

HD10.6 cells were maintained in maturation media for 7 days prior to stimulation. For stimulation experiments, complete media was replaced with Neurobasal Plus medium only for 1 hour in a 5% CO2 37°C incubator. Forskolin (25µM) or Phorbol 12-myristate 13-acetate (PMA) (1µM) (LC Lab, P-1680) was applied, and cells were returned to the incubator for 5 minutes. The stimulation time for in vitro experiments was optimized previously [26]. For western blotting, the medium was then aspirated entirely and replaced with 10% trichloroacetic acid (TCA) (Millipore, T0699) containing 20mM dithiothreitol (DTT) (Fisher Scientific, BP172-5) (TCA/DTT) to stop biochemical processes. Cells were incubated on ice for 20 minutes, then scraped in TCA/DTT, collected in Eppendorf tubes, and centrifuged for 15 min/15,000g to precipitate protein. TCA was aspirated completely, and pellets were washed in ice-cold acetone and then centrifuged for 5 min at 15,000g; this wash step was performed twice. Pellets were left to dry at room temperature in a fume hood for up to 20 minutes, solubilized in 1X Laemmli sample buffer (pH8.5), and shaken in an Eppendorf thermomixer at 25°C at 2000 rpm for 1 hour. Lysates were quantified (1.0 µl, in triplicate) with the EZQ protein quantification kit (ThermoFisher, R33200) and stored at −80°C for further analysis.

### Western blotting

SDS-PAGE was performed using Bolt bis-Tris 12% gels (Fisher Scientific, NW00122BOX) and Bolt MOPS SDS running buffer (Fisher Scientific, B0001). Proteins were transferred onto PVDF membranes (PB5240) using a power blotter system (Invitrogen, PB0013) set at 25 volts and 1.3 milliamperes for 10 minutes. PVDF membranes were probed with No-Stain protein labeling reagent (ThermoFisher, A44717) and imaged in the Cy3 channel before blocking for 1 hour at room temperature in 2.5% Bovine Serum Albumin (BSA) diluted in Tris-buffered saline (TBS) with 0.05% TWEEN-20 (TBST). PVDF membranes were incubated overnight on a rocker at 4°C with primary antibodies **(Table 2)** diluted in 0.5% BSA, 0.005% TWEEN-20 in TBS. After 24 hours, membranes were washed 3X10 minutes in 1X TBST buffer and incubated with donkey anti-rabbit Cy5 (Jackson ImmunoResearch, 711175152) or donkey anti-mouse Cy3 (Jackson ImmunoResearch, 715-165-150) diluted 1:2000 in 0.5% BSA in 1X TBS with 0.05% TWEEN-20 for 2 hours on a rocker at room temperature. Membranes were washed 3X10 minutes in TBST buffer and imaged wet using a Typhoon 9500 laser scanner (G.E. Healthcare Life Sciences). Band intensities were quantified using AutoQuant imaging software and normalized to total protein.

**Table 2:**
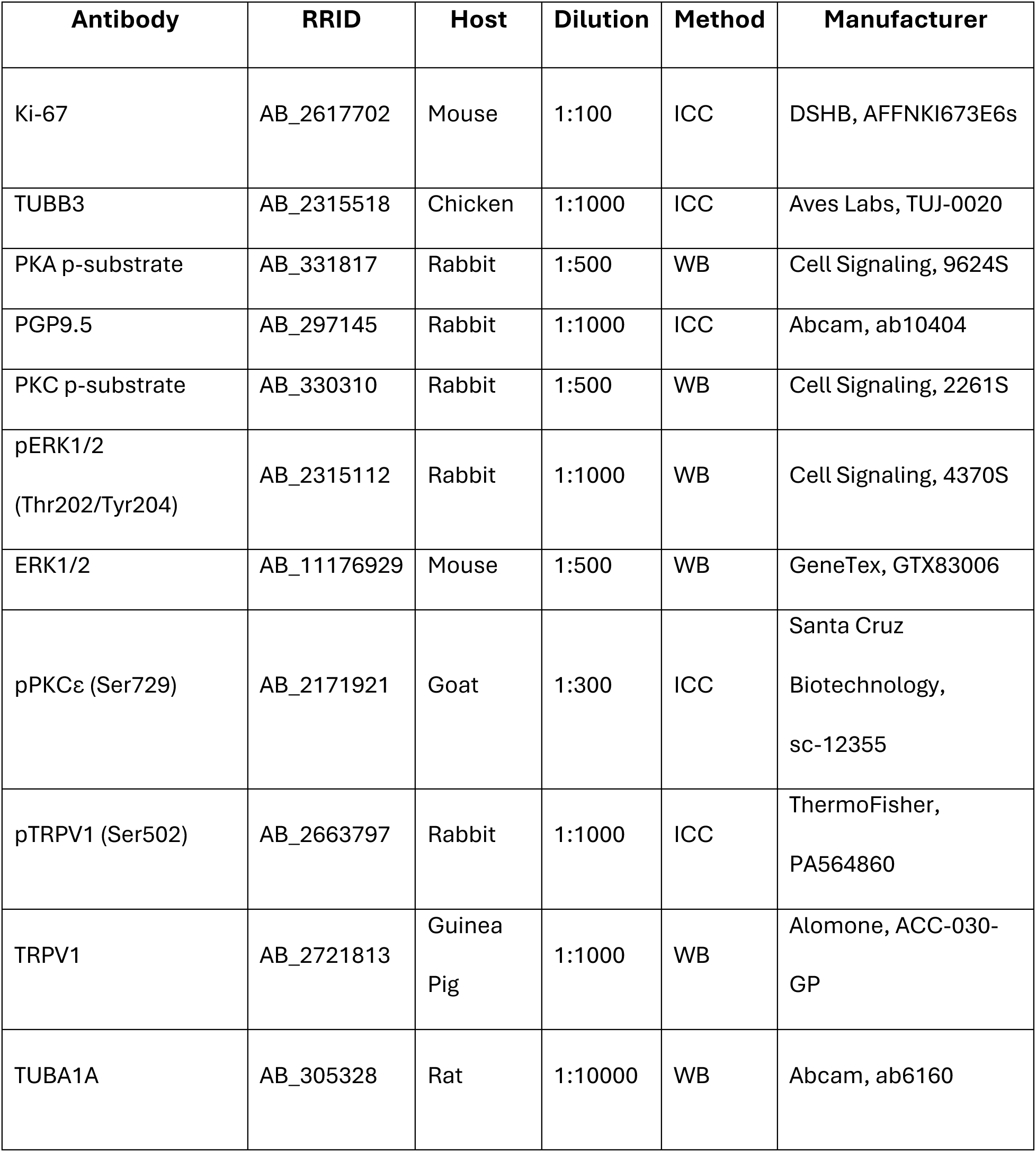
Primary antibodies. **ICC:** Immunocytochemistry, **WB:** western blot. **RRID:** Research Resource Identifiers.

### Immunocytochemistry (ICC)

Mature or proliferating HD10.6 cells were grown for 7 or 1 day, respectively, in 4-well EZ chamber slides (Millipore, PEZGS0416) coated with 5 µg/ml poly-D-lysine. For stimulation experiments, mature cells were pre-incubated in Neurobasal Plus medium for 1 hour at 5% CO2 37°C before stimulation with PMA (1µM) for 5 minutes. Cells were washed once in PBS (pH7.4), fixed in cold 4% paraformaldehyde for 20 minutes and washed 3 x 5 minutes in PBS before being incubated for 30 minutes in blocking buffer containing 2% normal donkey serum (Jackson Immunochemicals, 017000121), 0.1% Triton X-100 (Sigma, X100), and 0.05% sodium azide in PBS (Ricca Chemical, R7144800). Slides were incubated overnight at room temperature in primary antibodies (**Table 2**) diluted in the same blocking buffer. The following day, slides were washed 3X5 minutes in 1X PBS and incubated for 30 minutes at room temperature in secondary antibodies diluted 1:500 in blocking buffer. Slides were washed 3X5 minutes in 1X PBS and coverslipped using Fluoromount-G™ mounting medium with DAPI (Thermo, 00495952). Slides were stored at 4°C and imaged using a Leica Stellaris laser scanning confocal microscope. Image analysis was performed using Fiji version 2.14 for Mac OS [27].

### Statistics

All statistical comparisons were performed using Fiji, Microsoft Excel, or R programming language. Data are presented as the mean ± SEM or ± standard deviation (specified in caption). Experiments were replicated at least 4 times, as noted in the text. Significance is indicated in figures as *p<0.05, **p <0.01,***p< 0.001, or ****p<0.0001. Statistical details for the snRNA-seq and bulk RNA-seq analyses are described in the “Bioinformatics analysis” section.

## RESULTS

### HD10.6 cells efficiently mature into nociceptor-like cells

To examine the efficiency of HD10.6 cell maturation from proliferating cells to a neuron-like phenotype, we examined changes in cellular morphology and staining for neuronal markers β-III Tubulin (TUBB3) [58] and Ubiquitin C-Terminal Hydrolase L1 (PGP 9.5) [97], and a marker of dividing cells (Ki-67) [27]. Positive staining was consistently seen for both TUBB3 and PGP9.5 in proliferating as well as in mature cells (**Fig 1A**). Staining was absent in cells incubated without primary antibodies. In mature cells, staining for TUBB3 and PGP9.5 revealed a profuse net of neurite outgrowth from the cell bodies, which became spheroidal and more compact. In contrast, nuclear Ki-67 staining appeared in all proliferating cells but was eliminated in cultures grown in maturation media for 7 days. The mean cell body diameter for mature cells was 23.66± 0.62 µm (n=35), smaller than the mean diameter reported for a sample of slow-conducting cultured human DRG neurons (42.8± 0.8 µm) [15]. The results demonstrate highly efficient maturation of HD10.6 cells to a neuron-like morphology when exposed to maturation media and confirm that mature cultures are not contaminated with proliferating cells. The expression of TUBB3 and PGP9.5 in proliferating cells indicates that even proliferating cells display some phenotypic characteristics of neurons.

**Figure 1).**
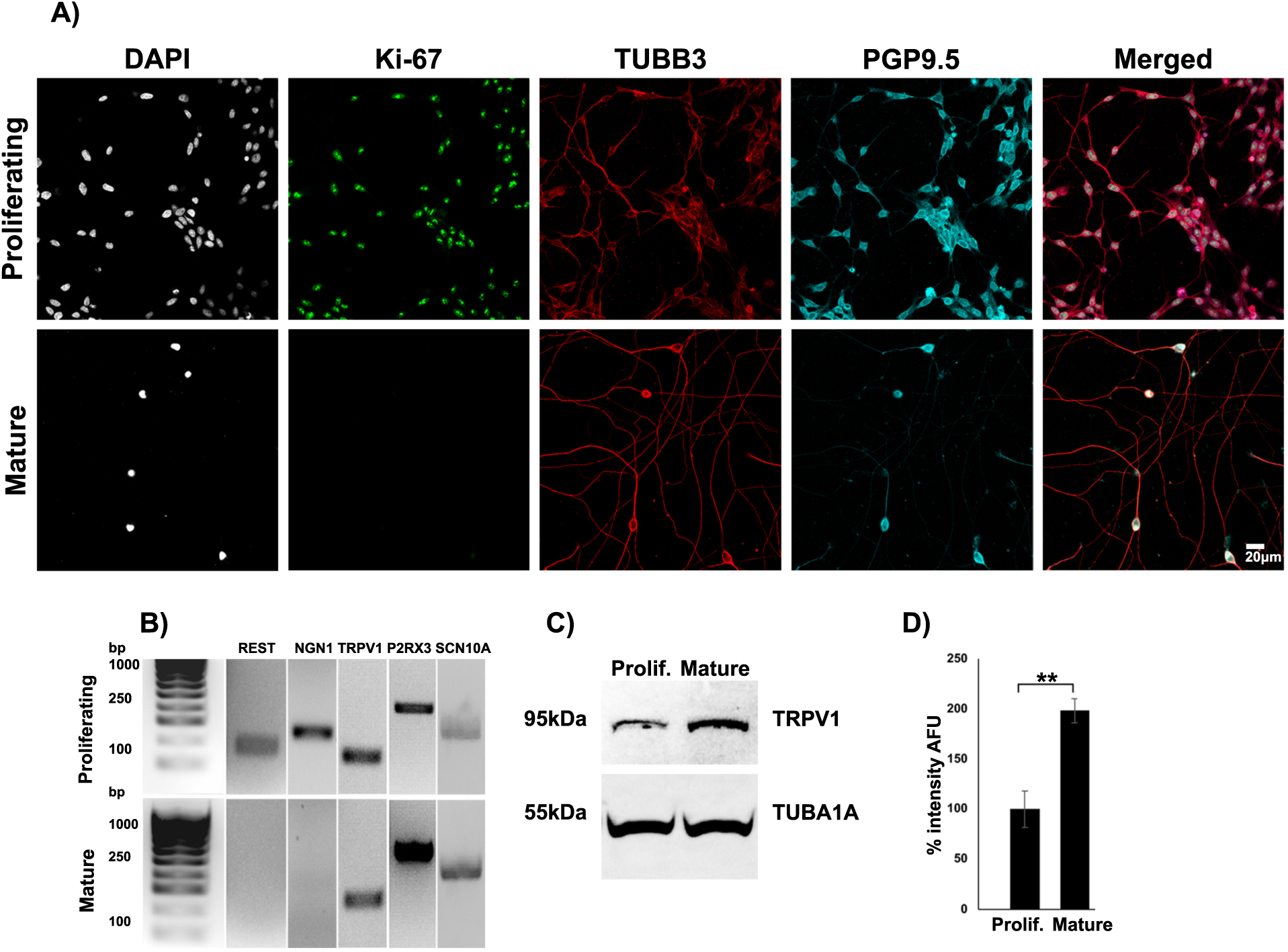
Maturation of HD10.6 cells is associated with changes in expression of sensory neuron markers. **A)** Immunocytochemistry (ICC) for neuronal markers beta-3 tubulin (TUBB3) and PGP 9.5 shows positive staining in mature and proliferating HD10.6 cells. Ki-67, a marker of dividing cells, labels proliferating cells but is absent in mature cells. The DNA stain 4ʹ,6-diamidino-2-phenylindole (DAPI) was used to identify the nuclei of all cells in cultures, regardless of phenotype. **B)** RT-PCR was used to test for the expression of several markers of sensory neurons in HD10.6 cells. Ion channels *TRPV1*, *SCN10A*, and *P2RX3* are expressed in both proliferating and mature cells. In contrast, transcription factors REST and NGN1 are expressed in proliferating cells but are downregulated in mature HD10.6 cells. **C)** Western blotting confirms the presence of TRPV1 and alpha-1A tubulin (TUBA1A) proteins in mature HD10.6 cells. **D**) Quantification of signal intensity in western blots for TRPV1 after normalization to TUBA1A demonstrates significant upregulation of TRPV1 protein in mature cells (n=4); ** = p<0.01.

For an initial evaluation of the HD10.6 cell gene expression profile, we used RT-PCR to examine the expression of several genes commonly associated with nociceptors, as well as several transcription factors associated with DRG neuron differentiation. Nociceptor markers included *SCN10A* (voltage-gated sodium channel Nav1.8), heat- and acid-gated ion channel *TRPV1*, and ATP-gated ion channel P2RX3. All three channel transcripts were detected in both proliferating and mature HD10.6 cells (**Fig 1B**).

Two transcription factors associated with sensory neuron development were examined: RE1 silencing transcription factor (*REST*, also referred to as NRSF) [41] and neurogenin 1 (*NEUROG1*, also referred to as NGN1). *REST* acts as a negative regulator of many neuron-specific genes in non-neuronal cells, as well as in neuroblasts prior to differentiation [80]. *NEUROG1* is a marker of DRG neuron progenitors that is required for the differentiation of most DRG nociceptors from neuroblasts [68]. Both genes were expressed in proliferating HD10.6 cells but were no longer detected after 7 days of exposure to maturation media (**Fig 1B**). The downregulation of these transcription factors during HD10.6 cell maturation is consistent with the process of DRG neuron differentiation that occurs *in vivo* [70; 81].

Finally, Western blotting was used to confirm the presence of *TRPV1* protein in both proliferating and mature HD10.6 cells and demonstrated a significant upregulation of TRPV1 protein in mature cells (**Fig 1C-D**).

### Differential gene expression between proliferating and mature HD10.6 cells

Once we confirmed that we could obtain consistent populations of mature HD10.6 cells, we performed bulk RNA-seq analysis of proliferating and mature cells to characterize the HD10.6 transcriptome (n = 5 replicates per condition). Principle component analysis (PCA) confirmed the consistency of cell culture replicates within conditions (**Fig S1A**). Of the 45,299 genes detected across all libraries, we observed 7,754 genes that were significantly upregulated in mature cells compared to proliferating cells (log_2_FC > 1 and Benjamini-Hochberg adjusted p-value < 0.05), while 6,581 genes were significantly downregulated (log_2_FC < −1 and Benjamini-Hochberg adjusted p-value < 0.05) (**Fig 2A**).

**Figure 2).**
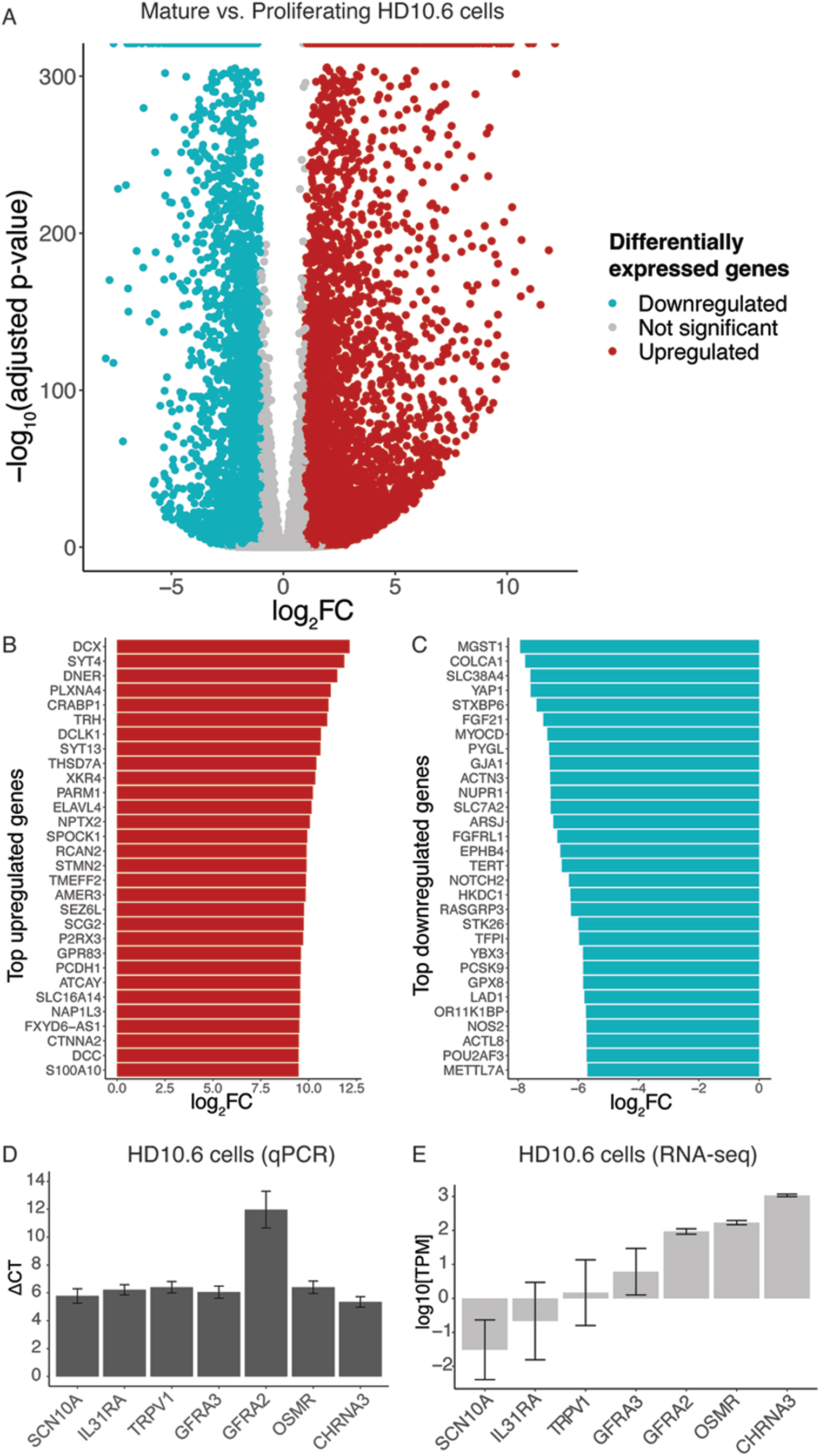
Gene expression changes during maturation of HD10.6 cells. **A**) Volcano plot illustrating the distribution of differentially expressed genes in mature HD10.6 cells compared to proliferating cells (log_2_FC>1, adjusted p-value<0.05). Top 30 upregulated **(B)** and downregulated **(C)** genes in HD10.6 cells compared to proliferating cells (log_2_FC>1, adjusted p-value<0.05). **D)** Expression levels of selected genes relevant to nociception/pruriception in mature HD10.6 cells, as measured by qPCR. The y-axis displays ΔCt values normalized to GAPDH (lower ΔCt values indicate higher levels of gene expression). **E)** Expression levels of the same genes from **(D)** in mature HD10.6 cells as determined by RNA-seq.

KEGG pathway analysis revealed that genes upregulated in mature cells were enriched for functions reflecting changes in retrograde endocannabinoid signaling, axon guidance, and neuroactive ligand-receptor interactions (**Fig S1B**). Many upregulated genes (**Fig 2B**) encode proteins that have been implicated in sensory neuron physiology, including the ion channel *SCN9A* [20; 22; 111], RNA-binding protein *ELAVL4* [45], neuronal migration protein *DCX* [5; 14; 47; 65], notch ligand *DNER* [24], membrane trafficking protein *SYT4* [100; 102; 106], and the β3 subunit of the gamma-aminobutyric acid (GABA-A) receptor *GABRB3* [62; 102; 106].

Genes downregulated in mature HD10.6 cells (**Fig 2C**) illustrate the extensive molecular and cellular changes these cells undergo during the transition to a mature state [63]. A number of genes involved in sensory neurogenesis in vivo are downregulated in mature HD10.6 cells, including *YAP1*, *MGST1*, and *ZFP36L1* [43; 48; 67; 89; 94]. Consistent with the RT-PCR results, we observed a downregulation of *REST* (log_2_FC = −4.58) and *NEUROG1* (log_2_FC = −1.63). Furthermore, several genes identified as targets for transcriptional suppression by REST are upregulated in mature HD10.6 cells (**Fig S2**), suggesting that the downregulation of REST may be responsible for the upregulation of a cohort of neuron-specific transcripts during maturation. These include multiple genes involved in neurotransmission and synaptic plasticity (e.g., *SCN2A, SCN8A, GRIN2A, BDNF,* and *ADCYAP1*) [21; 50; 82; 96; 101; 109].

To confirm the transcriptomics results, we performed qPCR for several genes of interest implicated in nociceptive/pruriceptive signaling: *CHRNA3, GFRA2, GFRA3, IL31RA, OSMR, SCN10A*, and *TRPV1*. All genes identified in HD10.6 cells by qPCR were also identified by RNA-seq (**Fig 2D-E**). However, *SCN10A* was lowly expressed in our RNA-seq data (average expression = 0.26 TPM), whereas it was robustly expressed by qPCR.

### Comparison of HD10.6 cells and iPSC-SNs to individual human DRG neuronal subtypes

A goal of this study was to compare the transcriptomic profile of mature HD10.6 cells to primary human nociceptor subtypes to determine whether features that distinguish human sensory neurons are retained in HD10.6 cells. We compared the transcriptomes from mature HD10.6 cells with the transcriptomes of human neuronal subtypes observed in the harmonized atlas recently published by Bhuiyan and colleagues [2]. We interpreted these correlations based on Schober et al. [63], who categorize correlations of 0.40–0.69 as “moderate” and 0.10-0.39 as “weak”, suggesting that values exceeding 0.40 are more likely to reflect meaningful transcriptomic correlations [75; 91]. Mature HD10.6 cells showed a moderate correlation (Spearman’s r > 0.40) with three nociceptive neuronal subtypes (Calca+Smr2, Mrgprd, and Sst; Spearman’s r = 0.6, 0.52, and 0.44, respectively) **(Fig. 3A)**. The transcriptomes of all iPSC-SNs screened also showed moderate correlations with individual DRG neuronal subtypes (Spearman’s r > 0.40 within the set of genes detected across all studies). A motor neuron iPSC line, included as a neuronal cell line control, also showed moderate correlations with multiple sensory neuron subtypes, in some cases exceeding those of any of the sensory neuron-differentiated iPSC lines for certain primary DRG subtypes. This suggests that while all screened cell lines share core neuronal molecular features, individual sensory neuron lines vary in how well they recapitulate primary sensory neuron gene expression profiles.

**Figure 3).**
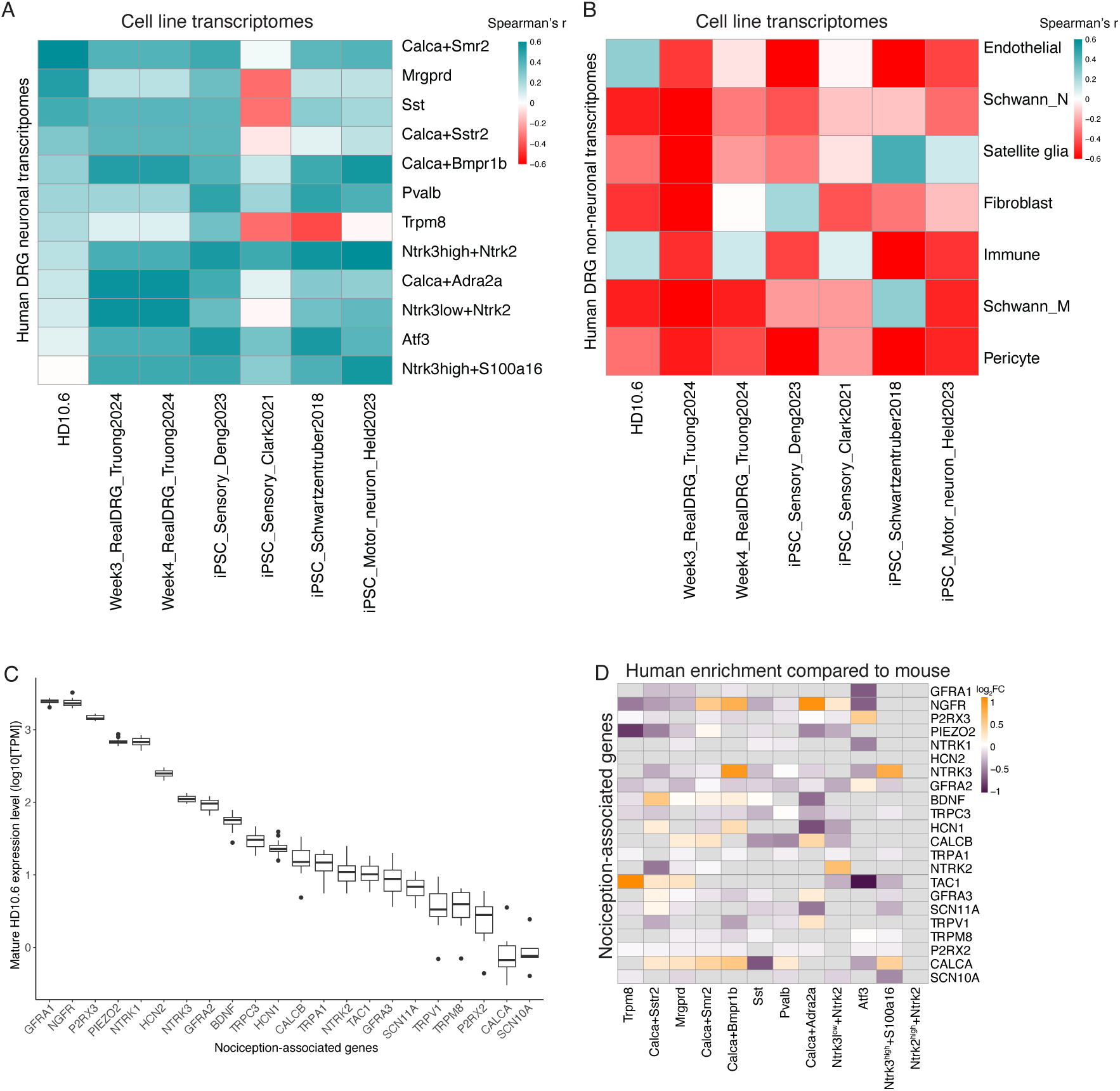
Correlation between transcriptomes from HD10.6 cells, iPSCs and human nociceptor subtypes. **A)** Correlation of transcriptomes from HD10.6 cells as well as recently developed iPSC-SNs and an iPSC motor neuron cell line to transcriptomes of human DRG neuron subtypes. The scale bar represents Spearman’s rank correlation coefficient (r). Only genes expressed across all datasets were used. **B)** Correlation of transcriptomes from mature HD10.6 cells and iPSC lines to transcriptomes of human DRG non-neuronal cell types. The scale bar represents Spearman’s rank correlation coefficient (r). Only genes expressed across all datasets were used. **C)** Expression levels of genes associated with nociceptive signaling in matureHD10.6 cells. The box plot displays mean log-normalized TPM values (y-axis; n=5). Error bars represent SEM. **D)** Heat map shows the enrichment of genes from **(C)** in human neuronal subtypes relative to the mouse in single nucleus/single-cell RNA-seq data. For each subtype, the heatmap displays log2FC enrichment for each gene in human nuclei compared to mouse cells/nuclei. Data were generated from [7].

HD10.6 cells exhibited negative to weak correlations (Spearman’s r < 0.40) with the non-neuronal DRG cell types (**Fig. 3B**). The only cell line that showed a negative correlation with all non-neuronal DRG subtypes was the RealDRG line differentiated for 3 weeks [104]. The other iPSC lines were negatively or weakly correlated with individual non-neuronal DRG cell types.

Notably, in comparing HD10.6-expressed genes across species in the harmonized atlas, we found 589 protein-coding genes expressed in both HD10.6 cells and human DRG neurons (>5 TPM) that do not have established orthologs in the mouse or rat genome **(Table S1)**. A literature search failed to reveal any investigations of these human genes for roles in nociceptive signaling. An additional 375 genes present in HD10.6 cells and human DRG neurons, but not in mouse or rat, are annotated as pseudo-genes (>5 TPM) **(Table S1)**. Taken together, these findings underscore the challenges of choosing a heterologous system for the study of neuronal signaling and support the use of HD10.6 cells as an *in vitro* model for human primary DRG neurons.

### HD10.6 cells express nociceptor marker genes

The RNA-seq analysis confirmed that HD10.6 cells express numerous transcripts implicated in nociceptive signaling (**Fig 3C**; see also [11; 37; 103; 118]). For example, HD10.6 cells express high levels of *P2RX3* and *PIEZO2*, which are highly expressed in both human and mouse primary neurons. They also express high levels of the NGF receptor tyrosine kinase *NTRK1* and the NTRK family co-receptor *NGFR*, which are broadly expressed in human and mouse nociceptors.

**Fig 3D** shows the extent to which the nociception-related genes evaluated in **Fig 3C** are differentially expressed in human compared to mouse. Several nociception-associated genes show substantial species differences in expression across sensory neuron subtypes. For example, in the Calca+Bmpr1b subtype, *NTRK3* is highly enriched in human compared to mouse. Moreover, in the Trpm8 subtype, TAC1 is highly enriched in human compared to mouse, whereas *PIEZO2* expression is substantially lower in the human subtype. For genes showing differences in expression between human and mouse, their expression in HD10.6 cells is largely consistent with human primary neurons. For example, the purinergic ion channel P2RX2 forms heteromers with P2RX3 in rodent C- and Aδ-nociceptive neurons [56], but there is little or no expression of P2RX2 in human DRG neurons, as noted previously [3], or in HD10.6 cells. These findings underscore the value of using human sensory neuron-like cells in investigating nociceptive signaling.

In contrast, we identified multiple genes for which HD10.6 cells diverge from human primary neurons. For example, the neuropeptide gene CALCA is expressed only at low levels in HD10.6 cells. HD10.6 cells also express relatively low levels of *TRPV1* mRNA compared to *P2RX3* and *PIEZO2*, although capsaicin-evoked currents have been demonstrated in these cells [37] and protein levels of TRPV1 are elevated in mature cells, compared to proliferating cells (see **Fig 1D)**. In addition, although we detected *SCN10A* by PCR, expression levels detected by RNA-seq were quite low (<1 TPM), although SCN11A (Nav1.9) levels were ∼6 TPM.

### Expression of ion channels in HD10.6 cells and iPSC lines compared to human sensory neuron subtypes

Transcripts for 562 distinct genes encoding ion channels were identified in HD10.6 cells (TPM >5) (**Fig S3**), many of which show subtype-specific expression in human DRG neurons (**Fig 4**). The most highly expressed channel transcript in HD10.6 cells, *CACNA1B*, encodes the pore-forming subunit of Cav2.2 N-type calcium channels. This protein is the target of ziconotide, an FDA-approved analgesic drug for refractory chronic pain derived from a cone snail toxin [72]. *CACNA1B* is expressed only at low levels in the iPSC lines compared to HD10.6 cells and human primary sensory neurons. Another ion channel that is highly expressed in HD10.6 cells and human primary sensory neurons but not in the iPSC lines is *SCN9A* (NaV1.7), a voltage-gated sodium channel that is essential for the manifestation of pain in mice and humans. Although variations in ion channel expression levels were observed across the different models, HD10.6 cells demonstrate comparable and in some cases greater similarities to primary human DRG neurons in ion channel gene expression than do the screened iPSC-SN lines.

**Figure 4).**
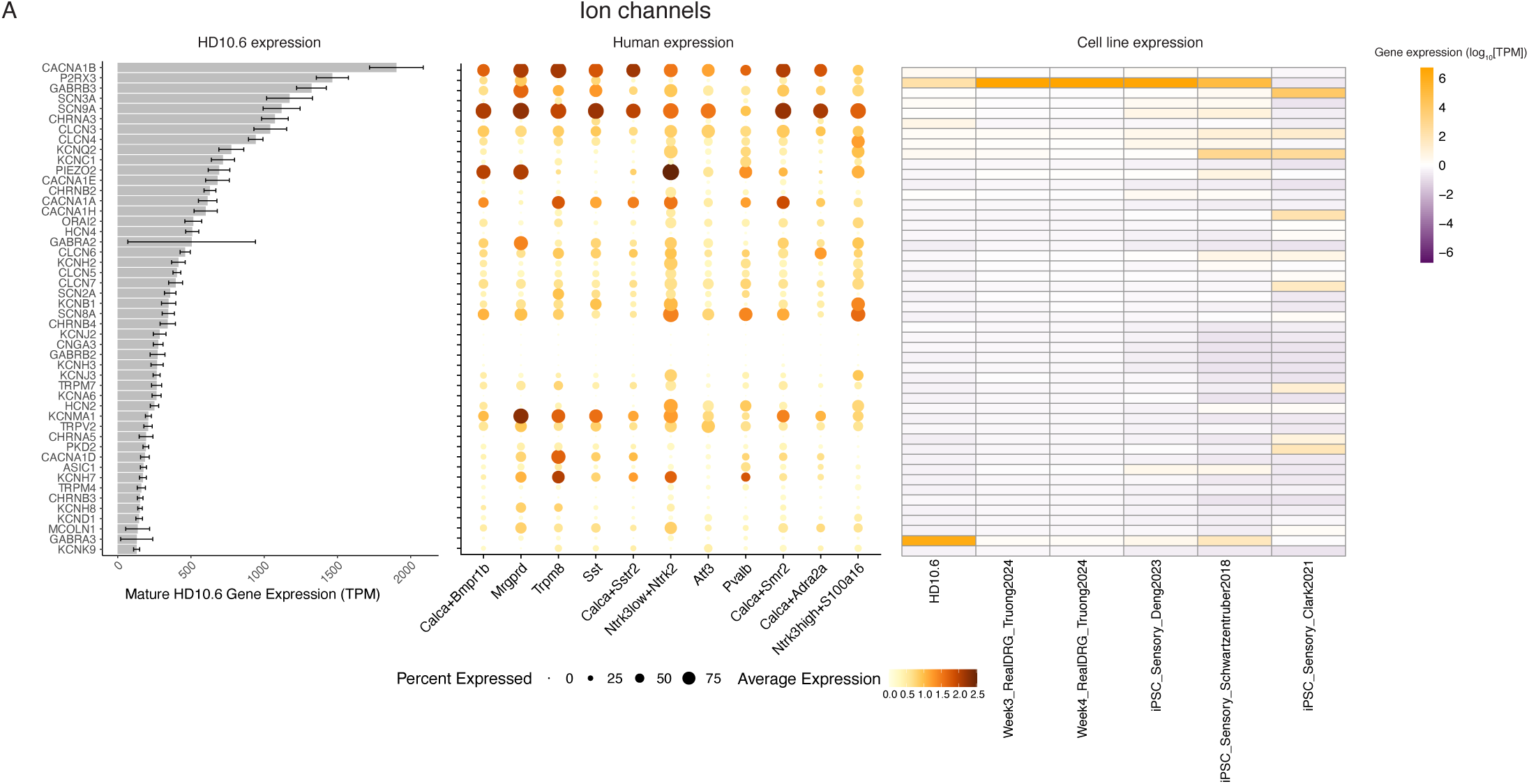
Expression of ion channels in HD10.6 cells, iPSC-SNs and human nociceptor subtypes. The expression of ion channels, which are enriched in human DRG neurons, in HD10.6 cells and human DRG neurons. **Left**: The bar plot displays average gene expression in mature HD10.6 cells. Error bars indicate standard deviation across HD10.6 samples (n=5). **Middle**: Dot plot of human neuronal subtype–specific gene expression. Dot size indicates the fraction of cells/nuclei expressing each gene, and color indicates the average log-normalized scaled expression of each gene. Data were generated from [7]. **Right**: Heatmap provides a comparison of log-normalized TPM levels between HD10.6 cells and recently developed iPSC-SNs.

One of the ion channel genes expressed in HD10.6 cells and human sensory neurons but not in mouse or rat is amiloride-sensitive sodium channel subunit delta (*SCNN1D)* [28] (**Table S1**); this transcript is expressed in multiple human nociceptor subtypes as well as in thinly myelinated low threshold mechanoreceptors (Aδ-LTMRs) and proprioceptors [7]. *SCNN1D* is reportedly activated by low pH [112] as well as icilin, a TRPM8 agonist [113]. As this gene is absent from the mouse genome, any contributions to sensory neuron excitability are unlikely to appear in the preclinical literature. The native expression of genes such as *SCNN1D* in HD10.6 cells provides a convenient *in vitro* model for analyzing their function and regulation.

### Expression of GPCRs by HD10.6 cells and iPSC lines compared to human sensory neuron subtypes

GPCRs are the most abundant family of transmembrane receptors and transduce extracellular stimuli into intracellular signaling processes [79]. They are targets for a large proportion of clinically used drugs, and numerous GPCRs have been implicated in nociceptor plasticity in rodent pain models [99]. In this study, we were particularly interested in the utility of HD10.6 cells for investigations of GPCR signaling mechanisms in human DRG neurons, and the extent to which these receptors can be investigated in other human cell lines. For this analysis, we generated a list of all non-olfactory mammalian GPCRs using the gpcrdb.org database [25]. Then we grouped GPCRs as coupling primarily to G**⍺**_i/o_, G**⍺**_s_, or G**⍺**_q/11_, as reported in the IUPHAR database, as described in the Methods. **Fig 5A-C** present comparisons of GPCR expression among HD10.6 cells, human primary DRG neurons, and the iPSC lines.

**Figure 5).**
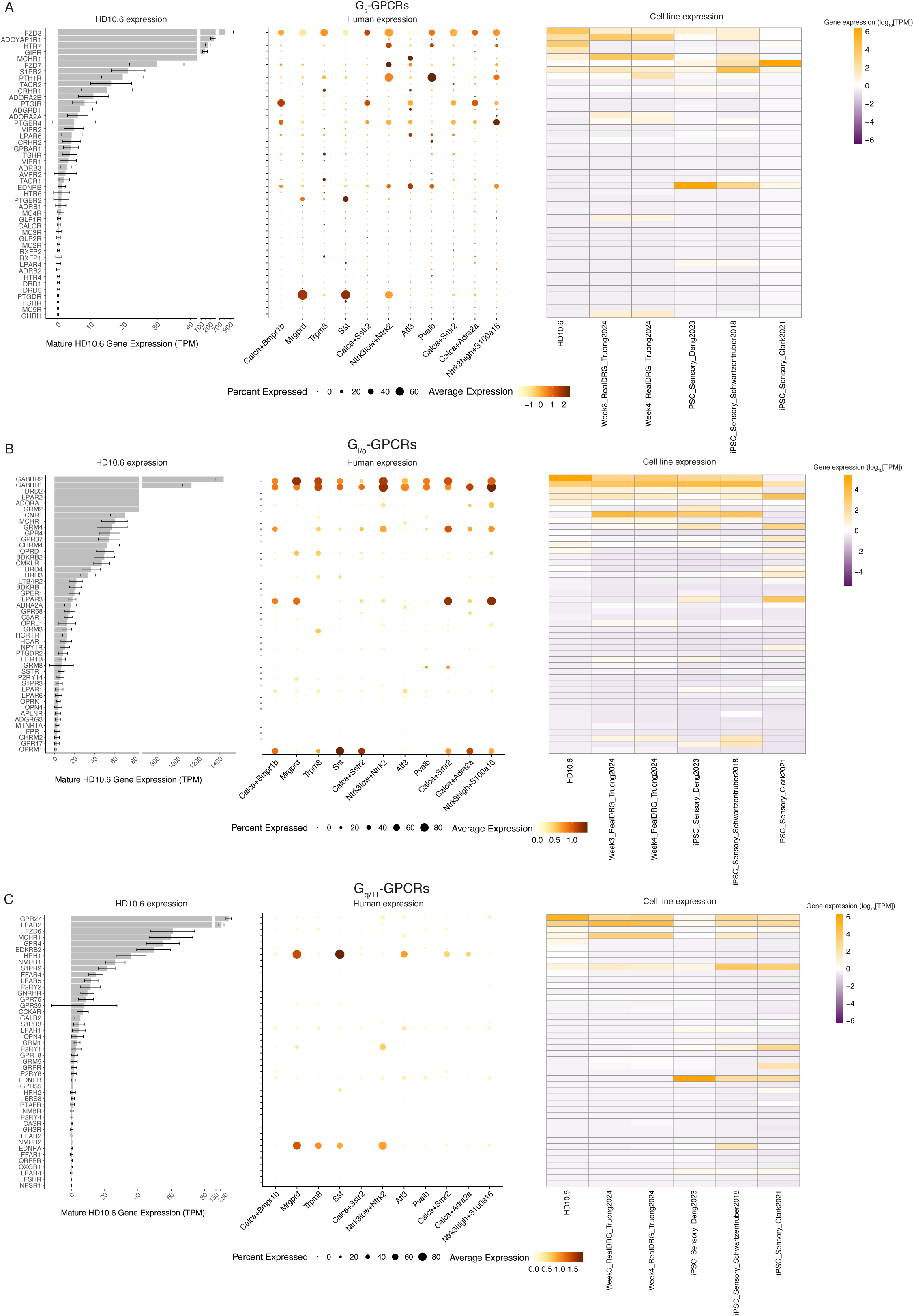
Comparison of GPCR expression in HD10.6 cells, iPSC-SNs and human nociceptor subtypes. Expression of receptors designated in the IUPHAR database as primarily G**⍺**_s_PCRs **(A)**, G**⍺**_i/o_PCRs **(B)**, or G**⍺**_q/11_PCRs **(C)** in mature HD10.6 cells and human DRG neurons. Receptors designated as coupling to multiple G proteins appear in multiple panels. **Left**: The bar plot displays average gene expression in HD10.6 cells. Error bars indicate standard deviation across HD10.6 samples. **Middle**: Dot plot of human neuron subtype–specific gene expression. Dot size indicates the fraction of cells/nuclei expressing each gene, and color indicates the average log-normalized scaled expression of each gene. Transcriptomics data were generated from [7]. **Right**: Heatmap displays log-normalized TPM expression values in mature HD10.6 cells and published iPSC lines.

Of the 415 non-olfactory GPCR genes screened, 303 were detected in mature HD10.6 cells (**Fig S4**). Of these, 195 receptors are categorized in the IUPHAR database as coupling to G protein types G⍺_i/o_, G⍺_s_, and/or G⍺_q/11_. One poorly characterized GPCR expressed in HD10.6 cells, the orphan receptor *GPR88*, is expressed at very low levels in naïve mouse and human neurons but is significantly upregulated in response to injury, and a synthetic agonist induced pain behavior in injured (but not naïve) mice [90]. Activation of GPR88 is associated with an increase in cAMP production, suggesting that it is coupled to G⍺_s_ [85]. Thus, it would be useful to study the regulation and function of this receptor in human models.

We identified 45 G⍺_s_PCRs in mature HD10.6 cells and analyzed their expression in primary human DRG neurons and iPSC lines (**Fig 5A**). This analysis revealed that some receptors are consistently expressed across almost all cell lines screened, such as *FZD3*, whereas others demonstrate distinct expression patterns in each cell line. For instance, the prostacyclin receptor (Ptgir) has been implicated in pro-inflammatory hyperalgesia in rodents [76; 84; 88], but its function has not been examined in human nociceptive signaling. We found that *PTGIR* is expressed in HD10.6 cells and several primary human nociceptor subtypes, however it is expressed only at very low levels in the iPSC-SNs that we screened. Another G⍺_s_PCR of interest, *P2RY11*, is expressed in both human sensory neurons and HD10.6 cells but is absent from the mouse and rat genomes (**Table S1**). *P2RY11* is the only identified G⍺_s_-coupled purinergic receptor [32; 49; 54]. Although multiple P2RY receptors have been implicated in nociceptive signaling in rodents, *P2RY11* has not been previously examined in human sensory neurons.

We detected 106 distinct G⍺_i/o_PCR transcripts in mature HD10.6 cells, making them the most diverse GPCR subset in these cells (**Fig 5B**). We compared the expression of G⍺_i/o_PCRs in HD10.6 cells, primary human DRG neurons, and iPSC lines. Despite many examples of receptors with similar expression levels across all models, such as GABBR1, HD10.6 cells express several G⍺_i/o_PCRs found in human primary neurons that are not detectable in the iPSC-SNs, including *GRM3, GRM8, LPAR6*, and *P2RY14*. In contrast, the µ opioid receptor OPRM1 is expressed only at low levels in HD10.6 cells and the iPSC-SNs, with the exception of the RealDRG cells.

We identified 47 G⍺_q/11_PCRs in mature HD10.6 cells (**Fig 5C**). Most of these receptors show similar expression patterns across all models. However, we identified several G⍺_q/11_PCRs in HD10.6 cells that are expressed in primary human DRGs but showed limited expression in the iPSC-SNs, including *HRH1* and *P2RY1*. The histamine receptor *HRH1*, which is significantly upregulated in mature HD10.6 cells and is the most highly expressed G⍺_q/11_PCR in human Sst and Mrgprd neuron subtypes, is detected only at low levels in the iPSC-SN lines. Pruriceptive *HRH1* actions are extensively documented [46]. It has been reported that phospholipase C beta 3 (Plcb3) is required for Hrh1 function in mouse, suggesting they may be physically coupled [34]. *PLCB3* and *HRH1* are also highly co-expressed in HD10.6 cells, providing an opportunity to investigate this relationship in human sensory neuron-like cells.

### Profiling activation of ERK1/2, PKA, and PKC in HD10.6 cells

We examined the profile of PKC phospho-substrate staining as a measure of PKC activity in HD10.6 cells, using an antibody targeting phospho-serine residues within the consensus recognition sequence for PKC (phospho-serine sites flanked by arginine or lysine at the −2 and +2 positions and a hydrophobic residue at the +1 position) [30]. As expected, direct activation of PKC isoforms with 1 μM PMA [66] for 5 minutes greatly increased PKC phospho-substrate staining (**Fig 6A**). PKC can also be activated downstream of G⍺_s_PCRs, through cAMP-responsive Rap guanine nucleotide exchange factors (*RAPGEF* family members), which signal primarily by activating Rap1 and Rap-dependent phospholipase C epsilon [23; 26; 40]. Consistent with this mechanism, PKC phospho-substrate staining was also induced, although to a lesser extent, in response to forskolin (FSK; 25 *μ*M, 5 min), an activator of adenylyl cyclases. The activation of PKC in response to cAMP production can be used as an assay for signaling by Rapgef family members downstream of G⍺_s_-coupled receptors [31].

**Figure 6).**
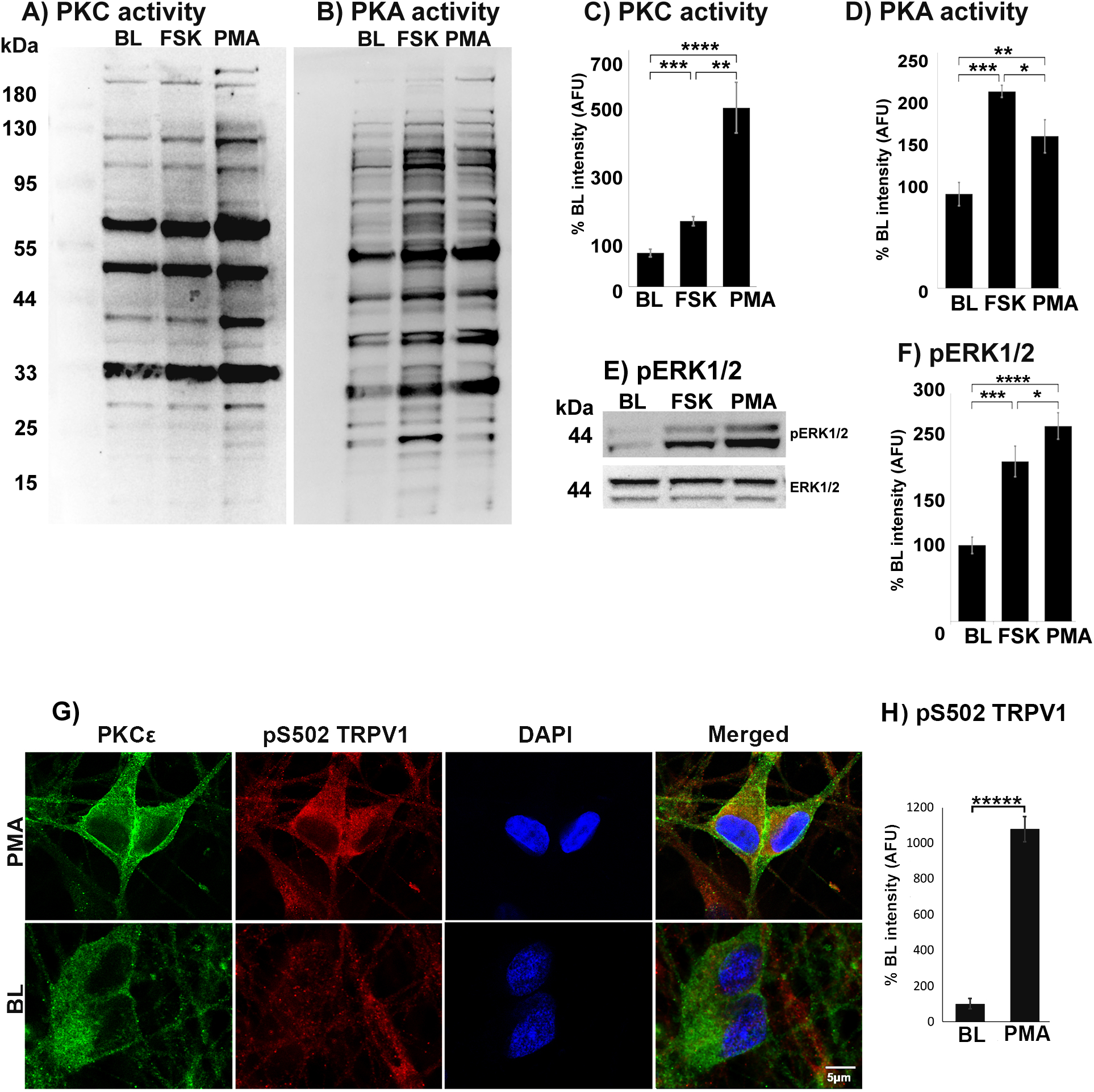
Measurement of PKA, PKC, and ERK signaling in HD10.6 cells. **A)** Western blot of PKC phospho-substrate staining in HD10.6 cells. BL: Baseline control. FSK: 25 µM forskolin. PMA: 1µM phorbol 12-myristate 13-acetate. PMA and FSK induced PKC activity to different extents, visualized as the increase in the intensity of PKC phospho-substrate staining across the entire lane. **B)** Western blot of PKA phospho-substrate staining in HD10.6 cells. FSK increased the intensity of PKA phospho-substrate staining. PMA induced a smaller but significant increase in PKA phospho-substrate staining. **C-D)** Quantification of overall staining intensity in **A-B**, normalized to total protein and compared to baseline (n=4). **E)** Western blot of pERK and total ERK signal in response to stimulation with either PMA or FSK. **F)** Quantification of the fluorescence intensity in (**E)** in arbitrary fluorescence units (AFU) of pERK, normalized to total ERK and compared to baseline (n=4). PMA, and to a lesser extent FSK, increased pERK signal intensity. **G)** ICC staining of mature HD10.6 cells demonstrates translocation of PKCε to the membrane and phosphorylation of TRPV1 in response to 5 minutes of stimulation with PMA. **H)** Quantification of the fluorescence intensity (in AFU) of phospho-TRPV1 signal intensity between BL control and stimulation with PMA. P values were calculated using t-test, p<0.05*, <0.01**, <0.001***, <0.0001****, <0.00001*****, n=4/condition.

To develop a profile of PKA phospho-substrate proteins in HD10.6 cells, we performed a similar assay using an antibody recognizing serine/threonine phosphorylation within the consensus recognition sequences for PKA (phospho-Ser/Thr residue with arginine at the −3 and −2 positions (RRXS*/T*)) [30]. We have previously validated this approach [30] and used it in this study to generate an electrophoretic band profile of PKA phospho-substrate proteins induced in response to FSK (25 *μ*M, 5 min) [93]. FSK significantly induced PKA phospho-substrate protein staining of diverse bands in protein samples from mature HD10.6 cells (**Fig 6B-D**).

GPCRs provide one of several major mechanisms for the activation of ERK1/2 in DRG neurons and other tissues [42]. G⍺_q/11_PCRs induce ERK1/2 phosphorylation (pERK1/2) principally through the activation of PKC [9]. G⍺_s_ activation of PKC through RAPGEFs can also lead to ERK activation [26]. In HD10.6 cells, pERK was induced in response to both FSK and PMA (**Fig 6E-F**). PMA evoked more robust phosphorylation, possibly due to its more direct and sustained activation of PKC [105]. In HD10.6 cells, pERK2 was more robust than pERK1 in response to either FSK or PMA. This result suggests the possibility of distinct roles for ERK1 and ERK2 downstream of GPCR signaling in human DRG neurons, whereas this difference in isoform activation was not observed in a report examining ERK1/2 activation in dissociated mouse DRG neurons [78]. However, this study did report that ERK2, but not ERK1, was required for the full manifestation of inflammatory nociceptor sensitization in mice [78].

Although multiple PKC isoforms have been implicated in nociceptor sensitization and hyperalgesia, PKCε is the most broadly implicated isoform in rodent models [52]. Among numerous substrates, TRPV1 is phosphorylated by PKCε at 2 sites (S502 and S800) in rodent DRG neurons, resulting in enhanced TRPV1 function and nociceptor hypersensitivity to heat [6; 69; 77; 108; 110; 121]. Like most PKC isoforms, PKCε is translocated to the plasma membrane upon activation, bringing it into contact with membrane-associated substrate proteins such as TRPV1. We used immunocytochemistry for PKCε to visualize the translocation of PKCε in HD10.6 cells. Application of PMA to HD10.6 cells for 5 minutes caused translocation of PKCε to the plasma membrane (**Fig 6G)**. Phospho-S502-TRPV1 staining was significantly increased in PMA-stimulated cells (**Fig 6H)**, providing evidence that TRPV1 is phosphorylated by PKC in human nociceptor-like cells, as previously described in rodents [6; 69; 77; 108; 110; 121].

## DISCUSSION

Animal models have been essential to developing our understanding of the molecular biology of nociceptive signaling and remain indispensable for some investigations, particularly behavioral and other in vivo analyses. Nevertheless, species differences can confound the translation of mechanisms identified in model organisms to human drug discovery efforts [51; 64; 119; 120]. The use of human neuronal cell lines, iPSCs, or isolated primary neurons for hypothesis generation and for validation of targets of interest identified in rodent studies can mitigate these translational risks, but validation against human expression data remains important. HD10.6 cells were generated 25 years ago [12] but have been used in only a small number of publications. In this study, we investigated the utility of HD10.6 cells for the study of human nociceptive signaling mechanisms by comparing the transcriptome of HD10.6 cells and several recently developed iPSC lines to the transcriptomes of human primary DRG neuron subtypes.

HD10.6 cells are derived from first-trimester human embryonic DRG [37]. DRG neurogenesis is largely complete by the end of the first trimester: central projections from the earliest-born (A-fiber) human DRG neurons into the spinal cord are initiated before week 8, and projections are extensive by week 15 [55]. For comparison, the peak of C-fiber neurogenesis in the rat occurs around embryonic day 14 [53], and central projections are abundant by embryonic day 17 [117]. Unlike iPSCs, HD10.6 cells are restricted to a neuron-like fate: they express some neuronal markers as proliferating cells and mature into neuron-like cells even in conditions optimized to induce glial differentiation [33]. When HD10.6 cells are cultured in maturation media, they undergo extensive transcriptional remodeling, including the upregulation of genes associated with nociception, pruriception, and thermosensation. Comparison of RNA-seq data from HD10.6 cells with published human DRG single nucleus RNA-seq data indicates that the HD10.6 cell transcriptomic profile is moderately correlated with 3 classes of putative nociceptors (Calca+Smr2, Mrgprd, Sst subtypes; Spearman’s *r* > 0.40 [91]), a stronger correlation than we found for several iPSC-SNs developed to study nociceptor physiology [12; 92]. HD10.6 cells show the strongest correlations with nociceptor subtypes rather than non-nociceptors or non-neuronal DRG cells. The moderate correlations of HD10.6 cells and iPSC-SN lines with multiple nociceptor subtypes suggests that these cell lines express numerous genes associated with a nociceptor-like identity, but do not uniquely reproduce any individual neuronal subtype. It should be noted that the strength of the transcriptomic correlations can be affected by differences in the resolution of different RNA-seq methodologies, such as RNA extraction protocols, sequencing platform, depth, and data normalization approaches. We attempted to control for as many of these variables as possible by limiting our analyses to genes detected across all datasets, though this comes at the cost of excluding numerous study-specific genes in the analysis, including nearly 30,000 genes excluded from the HD10.6 cell line dataset for this comparison. The majority of excluded genes were pseudogenes, long non-coding RNAs, and small non-coding RNAs. By restricting our analysis to genes common to all datasets, our comparisons minimize the impact of technical variations between datasets.

Despite the findings that HD10.6 cells and some iPSC-SNs correlate well with primary DRG neuron subtypes, examples of gene expression levels that diverge from those of primary neurons highlight the limitations of these models. Optimization of iPSC differentiation protocols is continuing to improve the consistency and translational efficacy of these models. These advances may also suggest adjustments to culture conditions for HD10.6 cells to more closely represent human nociceptor expression profiles.

HD10.6 cells express numerous ion channel and GPCR transcripts with unexplored potential as therapeutic targets. A notable finding from our analysis is the expression of a large cohort of genes in mature HD10.6 cells that are expressed in human sensory neurons but do not have established orthologs in mouse or rat. These genes may contribute to human pro- or anti-nociceptive mechanisms and cannot be investigated in native rodent models. A subset of these genes are annotated as pseudo-genes, however these may be worthy of additional evaluation since some pseudo-genes, such as GNRHR2, [17] have been subsequently documented to encode functional proteins [10]. The broad repertoire of GPCRs expressed in HD10.6 cells, including receptors yet to be investigated in sensory neurons such as *GPR107* and *GPR4* [60], offers a rich biochemical landscape for investigating metabotropic signaling pathways involved in sensory transduction and modulation.

There are several caveats related to our analysis of GPCRs. First, most studies cited by the IUPHAR measured inositol 1,4,5-trisphosphate (IP3) production to identify receptors coupled to G⍺_q/11_ [36]. However, the identification of *RAPGEF* family members as cAMP effectors [65] revealed a mechanism by which cAMP can activate phospholipase C, suggesting that some GPCRs identified as coupling to both G⍺_s_ and G⍺_q/11_ in databases such as gpcrdb.org [25] and PANTHER [71] may signal exclusively through G⍺_s_. Additionally, data from IUPHAR in the gpcrdb.org database is not yet updated to the latest version of the IUPHAR database [36], which includes updated information on GPCR coupling.

In **Table 3**, we compare the advantages and disadvantages of different *in vitro* models for human sensory neurons. Human primary DRG cultures are the closest to in vivo human neurons for low throughput analyses such as patch clamp electrophysiology to assess GPCR modulation of channel function [95]. Capabilities for manipulating human primary neurons in culture are rapidly evolving [57]. Rodent dissociated DRG cultures remain the most widely used *in vitro* model for ease of use and have greatly advanced our understanding of nociceptor physiology [59]. However, gene expression, protein function, and molecular networks may diverge from those of human [95]. Several rodent DRG-derived cell lines are available and provide the benefits of immortalized cells, but share the translational caveats of rodent neurons and may lack important human nociception-associated genes [115]. Non-neuronal human cell lines such as HEK and HeLa cells remain commonly used to express and characterize human ion channels [2; 13; 29] and other genes, yet these cells may lack neuron-specific binding partners, kinases, etc., that are critical for regulating protein function in the native human sensory neuron environment.

**Table 3.** Comparison of *in vitro* models of DRG neurons. Advantages and disadvantages of different models used for studying DRG neuron physiology.

| **Cells** | **Advantages** | **Disadvantages** |
| --- | --- | --- |
| **HD10.6 cells** | Human DRG-derived<br>Moderate correlation with human nociceptor subtype transcriptomes<br>Relatively simple, rapid 7-day protocol<br>Highly reproducible, restricted fate<br>Ease of genetic modification<br>Relatively inexpensive<br>Scalable; can be banked, disseminated | Does not fully reproduce primary nociceptor physiology<br>No option for patient genotype or history<br>Cannot be generated from patients with disease<br>Male only |
| **Human iPSCs** | Human-derived<br>Pluripotent: can be differentiated into many different subtypes<br>Moderate correlation with human nociceptor subtype transcriptomes<br>Potential access to source patient genotype/medical history<br>Option for both male and female cells<br>Ease of genetic modification<br>Potentially scalable; progenitor cells can be banked, disseminated | Differentiation can be time-consuming<br>Phenotype is highly sensitive to culture conditions<br>Does not fully reproduce primary nociceptor physiology<br>Commercially available cells typically cannot be propagated |
| **Human primary DRG neurons** | Most representative of in vivo human neurons<br>Potential access to donor genotype/medical history<br>Option for both male and female cells<br>Evolving protocols for in vitro manipulation | Requires access to donors<br>Not typically available from live patients<br>Not scalable, cannot be banked<br>Limited abundance<br>Quality inversely correlated with post-mortem interval<br>Heterogenous |
| **Rodent DRG neurons** | Relatively easy to obtain and culture<br>Correlated with human nociceptor subtypes<br>Reproducible, committed neuronal fate<br>Potential for in vivo manipulations<br>Flexible genetic background<br>Option for both male and female cells | Heterogeneous<br>Does not fully reproduce human physiology<br>Limited abundance (<~500,000 neurons per animal).<br>Cannot be banked, disseminated |

There are substantial benefits to using an immortalized cell line derived from human DRG as a complementary tool, particularly for high throughput and large volume biochemical and screening assays, including the ability to propagate large volumes of cells, to disseminate frozen aliquots for widespread use, and to readily transfect cells for experimental manipulations and functional imaging approaches. Our results visualizing phospho-substrate profiles for PKA and PKC, and the phosphorylation of ERK and TRPV1, provide initial demonstrations in human sensory neuron-like cells of well-established signaling pathways associated with nociceptor sensitization in rodents.

HD10.6 cells will not be appropriate for all in vitro investigations of nociceptive signaling; we describe some notable examples of HD10.6 cell transcript expression that diverges from the RNA-seq profiles for putative primary nociceptor subtypes, such as the low levels of neuropeptide *CALCA,* opioid receptor *OPRM1*, and the previously reported absence of TTX-resistant sodium currents [37]. While we detected transcripts for *SCN10A* by qPCR and for *SCN11A* by RNA-seq, functional protein may be absent or may fail to be inserted into the plasma membrane. Further investigation is needed to determine whether SCN10A and/or SCN11A expression can be enhanced by altering the culture conditions or translational control factors. These variations from human DRG gene expression profiles demonstrate the importance of referencing ‘omics data for validation of translational relevance to human primary neurons when using *in vitro* models, whether cell lines or rodent neurons. It is our hope that the transcriptomic analysis reported here will provide a useful resource for determining when HD10.6 cells are an appropriate choice for investigations of nociceptive signaling mechanisms and the development of high-throughput screening assays.

A point for consideration is that the maturation media used in this study was based on previous publications with HD10.6 cells, and the application of neurotrophic factors likely impacted the HD10.6 gene expression profile [1]. Additionally, caution must be taken when interpreting transcriptomics data because translational regulatory mechanisms, including several RNA-binding proteins identified here in both human neurons and HD10.6 cells, can cause protein levels to diverge from mRNA levels [86]. Finally, our correlation of cell line transcriptomes to human DRG neuron subtype-specific transcriptomes is limited by the quality of the human single nucleus RNA-seq data. As additional, more deeply sequenced, and higher quality human single cell RNA-seq data are generated, the transcriptomic similarity between HD10.6 cells and each neuronal subtype will likely shift. The most accurate comparison would be between single cell RNAseq data from both primary sensory neurons and HD10.6 cells, which are not yet available for HD10.6 cells.

Ultimately, choosing the most appropriate model for studying human nociceptive signaling will depend on the specific research question to be addressed. Overall, the relatively strong expression of many nociception-associated genes in HD10.6 cells and the native expression of many human genes absent in mouse support the use of HD10.6 cells as an accessible and reproducible model for the investigation of human sensory neuron signaling mechanisms.

## ACKNOWLEDGEMENTS

We thank Dr. Victor Hsia (College of Pharmacy, University of Maryland, Eastern Shore), for kindly providing HD10.6 cells, and we thank Emily Addleson for technical support and optimization of HD10.6 cell culture. We thank Peter Caradonna, the manager of the UNE Center for Pain Research Histology & Imaging Core (supported by P30GM145497), for assistance with fluorescence imaging of cell culture slides. The authors would like to thank members of the Molliver lab and Renthal lab for their guidance throughout the study. This project was supported by Burroughs Wellcome Fund (W.R.), Rita Allen Foundation (W.R.), Migraine Research Foundation (W.R.), and the National Institute of Neurological Disorders and Stroke [U19NS130617 (W.R.), R01NS119476 (W.R.), R01NS109936 (DCM)]. W.R. also receives support from the National Institute of Drug Abuse (DP1DA054343), National Eye Institute (U01EY034709), Teva Pharmaceuticals, Pfizer, BWH Women’s Brain Initiative, BWH Neurotechnology Studio, and MGB Gene and Cell Therapy Institute. W.R. is a consultant for Grunenthal and Eli Lilly. The RNA seq analysis reported in this publication was supported by an NIH Institutional Development Award (IDeA) Maine INBRE core access award under grant number P20GM103423 (ZA).

## Data Availability

The HD10.6 bulk RNA-seq dataset generated in this study has been deposited in the Gene Expression Omnibus (GEO) repository under accession number PRJNA1156623. The processed data are provided in **Table S1**. The human reference genome GRCh37 used in this study is publicly available from the National Human Genome Research Institute (NHGRI). As noted in the text, all analyses are based on previously published code and software.

