## Supplementary Figures and Legends for "A Transcriptomic Comparison of the HD10.6 Human Sensory Neuron-Derived Cell Line with Primary and iPSC Sensory Neurons"

**LIST OF SUPPLEMENTARY FILES**

Figure S1. Consistency of HD10.6 cell maturation and gene expression.

Figure S2. The expression levels of genes that are negatively regulated by REST.

Figure S3. The expression levels of ion channels in mature HD10.6 cells.

Figure S4. The expression levels of GPCRs in mature HD10.6 cells.

Table S1. Gene expression values from proliferating and mature HD10.6 cells.

### Figure S1

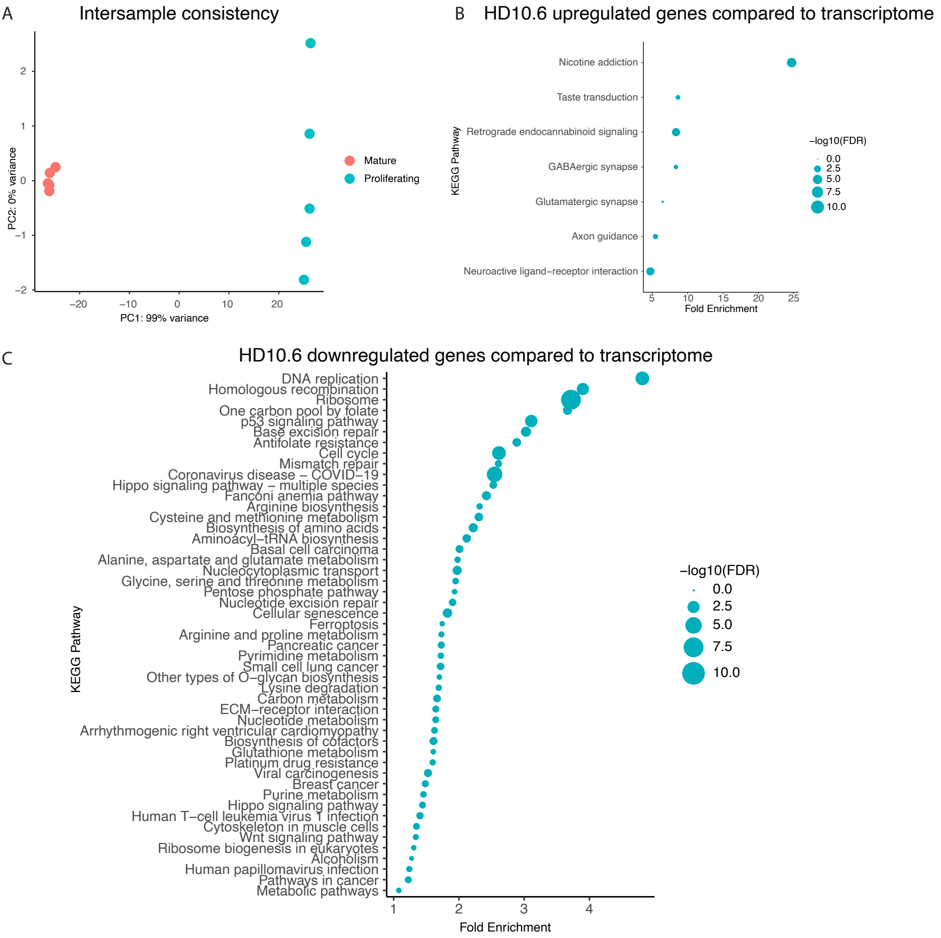
A
Intersample consistency
PC2: 0% variance
PC1: 99% variance
Mature
Proliferating
B
HD10.6 upregulated genes compared to transcriptome
KEGG Pathway
Nicotine addiction
Taste transduction
Retrograde endocannabinoid signaling
GABAergic synapse
Glutamatergic synapse
Axon guidance
Neuroactive ligand−receptor interaction
Fold Enrichment
−log10(FDR)
C
HD10.6 downregulated genes compared to transcriptome
KEGG Pathway
DNA replication
Homologous recombination
Ribosome
One carbon pool by folate
p53 signaling pathway
Base excision repair
Antifolate resistance
Cell cycle
Mismatch repair
Coronavirus disease − COVID−19
Hippo signaling pathway − multiple species
Fanconi anemia pathway
Arginine biosynthesis
Cysteine and methionine metabolism
Biosynthesis of amino acids
Aminoacyl−tRNA biosynthesis
Basal cell carcinoma
Alanine, aspartate and glutamate metabolism
Nucleocytoplasmic transport
Glycine, serine and threonine metabolism
Pentose phosphate pathway
Nucleotide excision repair
Cellular senescence
Ferroptosis
Arginine and proline metabolism
Pancreatic cancer
Pyrimidine metabolism
Small cell lung cancer
Other types of O−glycan biosynthesis
Lysine degradation
Carbon metabolism
ECM−receptor interaction
Nucleotide metabolism
Arrhythmogenic right ventricular cardiomyopathy
Biosynthesis of cofactors
Glutathione metabolism
Platinum drug resistance
Viral carcinogenesis
Breast cancer
Purine metabolism
Hippo signaling pathway
Human T−cell leukemia virus 1 infection
Cytoskeleton in muscle cells
Wnt signaling pathway
Ribosome biogenesis in eukaryotes
Alcoholism
Human papillomavirus infection
Pathways in cancer
Metabolic pathways
Fold Enrichment
−log10(FDR)

**Figure S1. Consistency of HD10.6 cell maturation and gene expression**

**A)** PCA plot illustrates the highly divergent molecular signatures between the proliferating and mature cell transcriptomes. Cell culture replicates are highly consistent within each condition. **B)** KEGG pathway analysis of genes upregulated (log2FC > 1, adjusted p-value < 0.05) in mature HD10.6 cells compared to proliferating cells. The size of the dot indicates -log10 of the false discovery rate (-log10(FDR)). **C)** KEGG pathway analysis of genes downregulated (log2FC > 1, adjusted p-value < 0.05) in mature HD10.6 cells compared to proliferating cells. The background is the list of all other expressed genes identified. The size of the dot indicates -log10(FDR).

### Figure S2

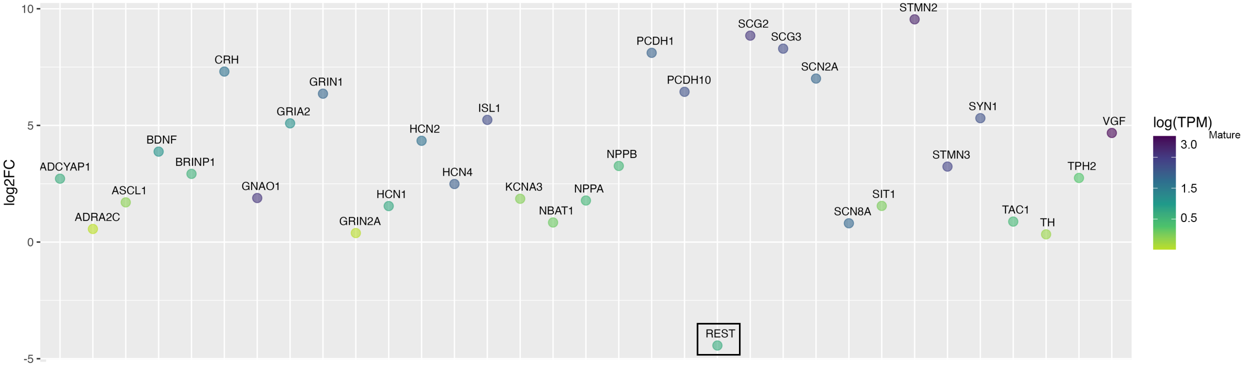

log2FC
10
5
0
-5
ADCYAP1
ADRA2C
ASCL1
BDNF
BRINP1
CRH
GNAO1
GRIA2
GRIN1
GRIN2A
HCN1
HCN2
HCN4
ISL1
KCNA3
NBAT1
NPPA
NPPB
PCDH1
PCDH10
REST
SCG2
SCG3
SCN2A
SCN8A
SIT1
STMN2
STMN3
SYN1
TAC1
TH
TPH2
VGF
log(TPM)
Mature
3.0
1.5
0.5

**Figure S2. The expression levels of genes that are negatively regulated by REST.** The left axis of the plot represents the log2 fold change in mean expression levels between proliferating and mature cells. The color gradient indicates the log10 expression level in mature cells, with purple indicating high TPM (transcripts per million) and yellow indicating low TPM. These genes were previously reported as REST-regulated genes [44]. The extent of the downregulation of REST in mature cells is highlighted with a black box.

**Figure S3**

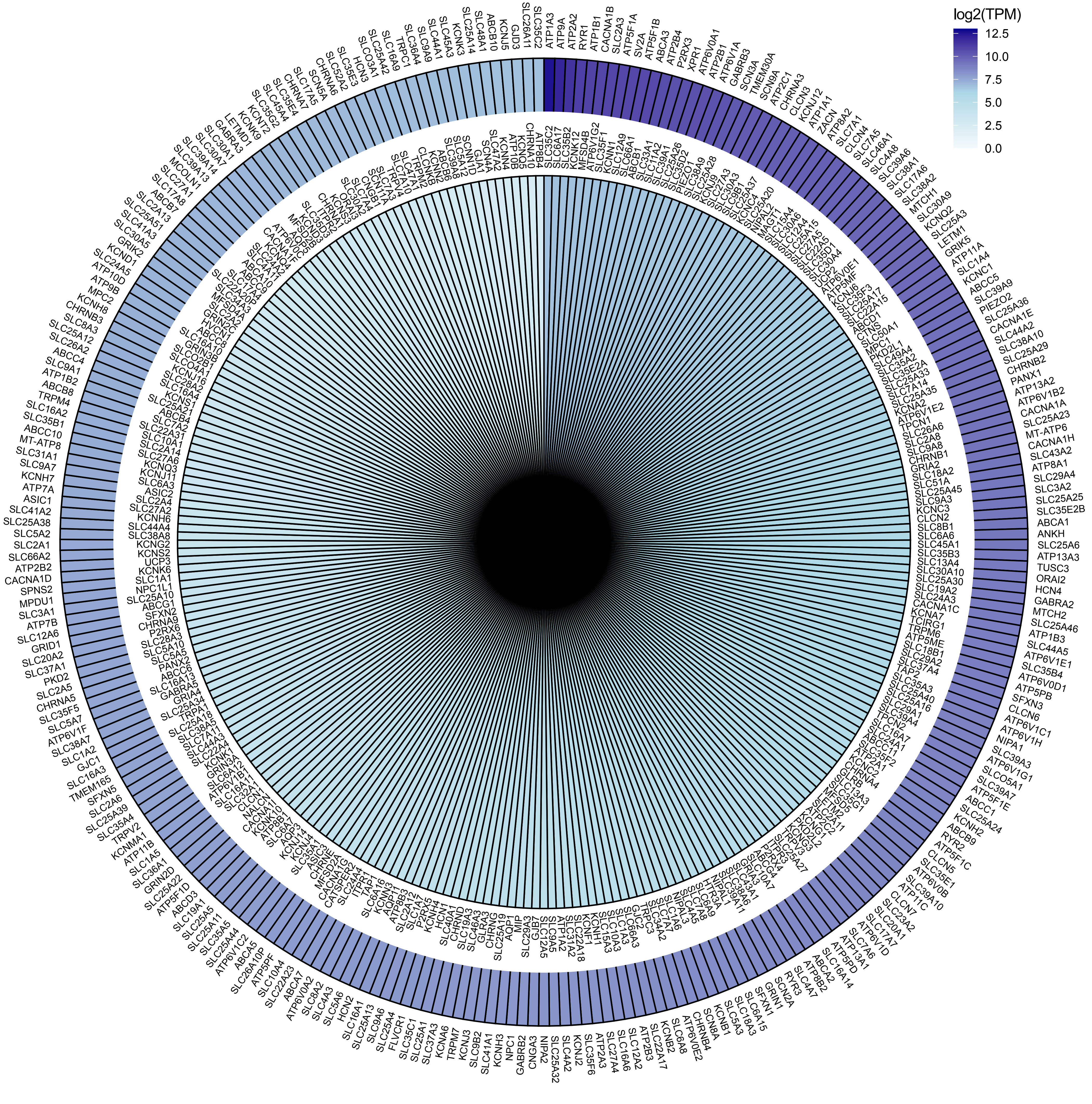

**Figure S3. The expression levels of ion channels in mature HD10.6 cells.** The color gradient of the radial bars on the plot represents log2(TPM) of the expression levels of all identified ion channels in mature HD10.6 cells, based on the PANTHER gene ontology designation.

**Figure S4**

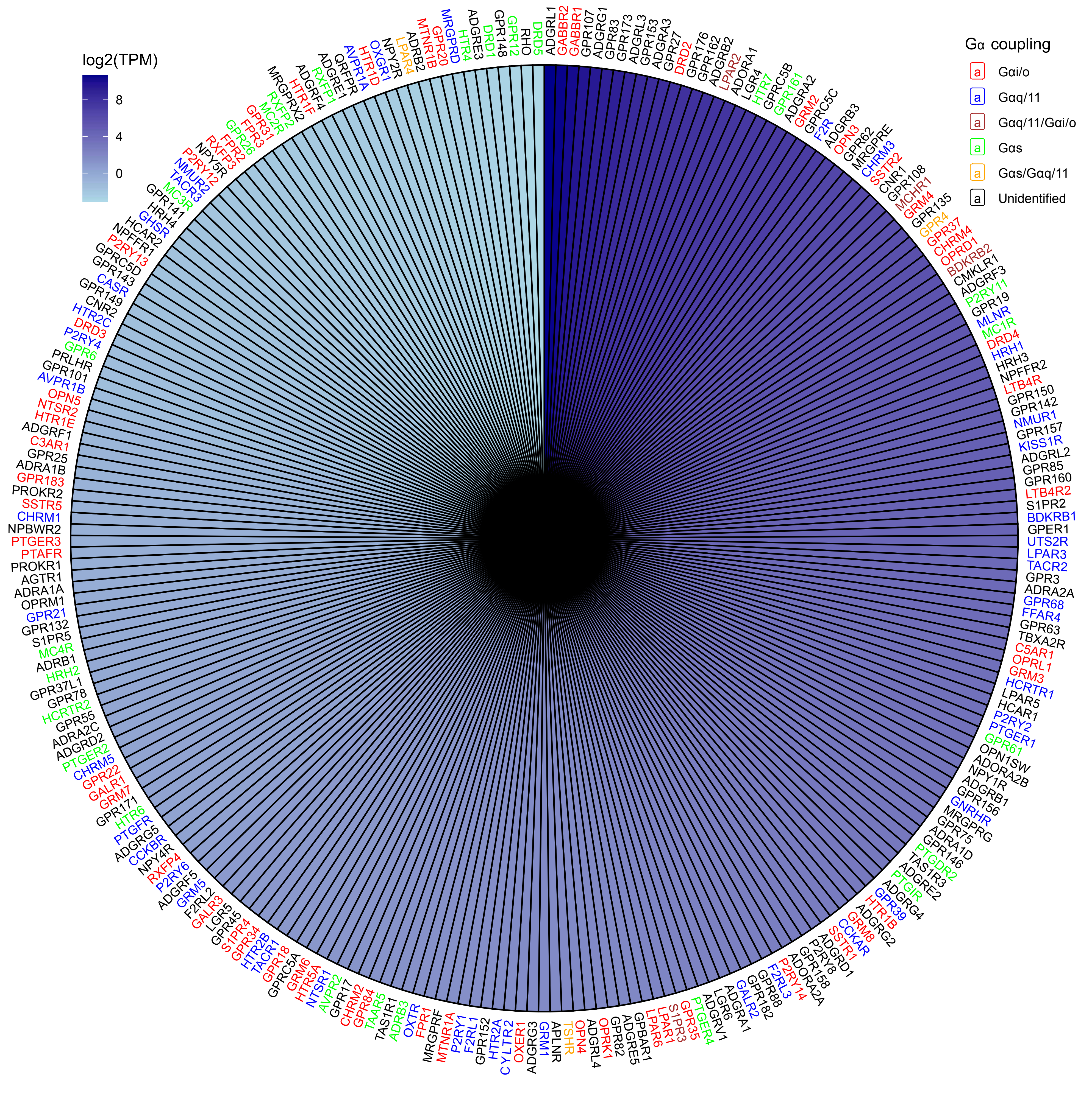

log2(TPM)
8
4
0
Gα coupling
a Gαi/o
a Gαq/11
a Gαq/11/Gαi/o
a Gαs
a Gαs/Gαq/11
a Unidentified

**Figure S4. The expression levels of GPCRs in mature HD10.6 cells.** This figure illustrates the expression levels of all non-olfactory GPCR genes identified in mature HD10.6 cell samples, identified as described in the Methods. The color gradient of the radial bars on the plot represents log2(TPM) of the expression levels. Gene names are color-coded to indicate putative Gα subunit coupling, based on the IUPHAR database [36]: $G\alpha_s$ in green, $G\alpha_{i/o}$ in red, and $G\alpha_{q/11}$ in blue. Promiscuous coupling is indicated with mixed colors: $G\alpha_s/G\alpha_{q/11}$ in orange, $G\alpha_s/G\alpha_{i/o}$ in yellow, and $G\alpha_{q/11}/G\alpha_{i/o}$ in brown. Coupling for receptors in black is undefined in the IUPHAR database.

**Table S1**. **Gene expression values in proliferating and mature HD10.6 cells**. The average expression levels (TPM) for all genes expressed in proliferating and/or mature HD10.6 cells. Table Tabs: **all genes**, contains all genes expressed in HD10.6 cells, regardless of expression level. HD10.6 genes > 5: Contains all genes expressed in mature HD10.6 cells with TPM ≥ 5. **Gs**, contains all G$\alpha_s$-coupled GPCRs expressed in mature HD10.6 cells. **Gi**, contains all G$\alpha_{i/o}$-coupled GPCRs expressed in mature HD10.6 cells. **Gq**, contains all G$\alpha_{q/11}$-coupled GPCRs expressed in mature HD10.6 cells. **Ion channels**, contains all ion channels expressed in mature HD10.6 cells. **Human genes**, contains genes expressed in human DRG neurons and HD10.6 cells that are absent in the mouse and rat genomes. **Human pseudogenes**, contains pseudogenes expressed in human DRG neurons and HD10.6 cells that are absent in the mouse and rat genomes. The expression levels of genes expressed in HD10.6 cells are present in human primary DRG neurons but not identified in the mouse genome. All listed genes were tested against the mouse genome obtained from Ensembl to validate absence. Genes were also loaded into the BioGPS.org tool to search for orthologs with different names in mouse or rat; genes with orthologs in mouse or rat were excluded. The total expression across primary human neuronal subtypes was summed, and genes with expression <5TPM were filtered out.
